# Ruxolitinib partially reverses functional NK cell deficiency in patients with *STAT1* gain-of-function mutations

**DOI:** 10.1101/157271

**Authors:** Alexander Vargas-Hernandez, Emily M. Mace, Ofer Zimmerman, Christa S. Zerbe, Alexandra F. Freeman, Sergio Rosenzweig, Jennifer W. Leiding, Troy Torgerson, Matthew C. Altman, Edith Schussler, Charlotte Cunningham-Rundles, Ivan K. Chinn, Imelda C. Hanson, Nicholas L. Rider, Steven M. Holland, Jordan S. Orange, Lisa R. Forbes

## Abstract

**Background:** Natural Killer (NK) cells are critical innate effector cells whose development is dependent on the JAK-STAT pathway. NK deficiency can result in severe or refractory viral infections. Patients with Signal Transducer and Activator of Transcription (STAT)1 gain of function (GOF) mutations have increased viral susceptibility.

**Objective:** We sought to investigate NK cell function in STAT1 GOF patients. Methods: NK cell phenotype and function were determined in 16 STAT1 GOF patients.

**Methods:** NK cell phenotype and function were determined in 16 STAT1 GOF patients.NK cell lines expressing patient mutations were generated with CRISPR-Cas9 mediated gene editing. STAT1 GOF NK cells were treated in vitro with ruxolitinib.

**Results:** Peripheral blood NK cells from of STAT1 GOF patients had impaired terminal maturation. Specifically, patients with *STAT1 GOF* mutations have immature CD56^dim^ NK cells with decreased expression of CD16, perforin, CD57 and impaired cytolytic function. STAT1 phosphorylation was elevated but STAT5 was aberrantly phosphorylated in response to IL-2 stimulation. Upstream inhibition of STAT signaling with the small molecule JAK1/2 inhibitor ruxolitinib *in vitro* and *in vivo* restored perforin expression in CD56^dim^ NK cells and partially restored NK cell cytotoxic function.

**Conclusions:** Properly regulated STAT1 signaling is critical for NK cell maturation and function. Modulation of elevated STAT1 phosphorylation with ruxolitinib is an important option for therapeutic intervention in patients with *STAT1 GOF* mutations.

## INTRODUCTION

Natural killer (NK) cells account for approximately 10-15% of all circulating lymphocytes^1^ and are important early effectors in the innate immune response to a variety of viral infections.^2^ Within peripheral blood, NK cells comprise two phenotypic and functional subsets.^3^ CD56^dim^ NK cells are considered to be terminally mature and are the primary mediators of contact-dependent lysis of target cells,^1, 3^ whereas the CD56^bright^ NK cells contain less mature NK cell subsets are potent producers of cytokines.^4, 5^

NK cells are derived from CD34^+^ precursors,^3, 6^ and their development can be stratified to 5 stages,^7^ characterized by specific surface receptors and proliferative and functional capacities.^8, 9^ CD56^bright^ NK cells are the minor subset (~10%) of NK cells within peripheral blood.^5^ They are characterized by low expression of perforin and high expression of CD94-NKG2A receptors,^7^ and a subset of cells retain CD117 (c-Kit) expression.^7, 10^ NKG2D, NKp46, CD62L and detectable CD122 are highly expressed.^10-12^ These cells are sources of cytokines reflected by high levels of interferon (IFN)-γ and granulocyte monocyte-colony stimulating factor (GM-CSF).^1, 7-10, 13-18^. A functional and phenotypic intermediate populations between CD56^bright^ and CD56^dim^ have also been described in healthy donors.^9, 11, 19^ NK cell terminal maturation is defined by up-regulation of perforin, CD16, CD57, CD8 and NKp46, and down-regulation of CD94.^7^, ^11,15-17^ NK cell development and homeostasis requires interleukin (IL)-15, ^20, 21^ and in both mouse and human systems NK cells do not develop in the absence of IL-15.^25^

Classical natural killer deficiency (cNKD) is characterized by the absence of both NK cells and their cytotoxic function,^23-27^ whereas functional NK cell deficiency (fNKD) is characterized by normal frequencies of NK cells in peripheral blood with decreased function.^28, 29^ There are several immune deficiency diseases that affect NK cell development and/or function. ^29^ Patients with NKD have increased susceptibility to viral infections, including herpesviruses, varicella zoster, herpes simplex, cytomegalovirus, and human papilloma virus.^23-27, 30^ Severe herpes viral infection with decreased NK cell natural cytotoxicity has been reported in patients with loss-of-function mutations in signal transducer and activator of transcription-(STAT) 1,^31^ and recently in 8 patients with STAT1 gain of function (GOF) mutations.^32^

The STAT family includes seven members: STAT1-4, STAT5a and b, and STAT6.^33^ Activation through intracellular domains of cytokine receptors, including those for IFN-α/γ, IL-2, IL-4, IL-15, IL-21, and IL-6^34-36^ leads to association with Janus kinase (JAK) family members and recruitment and phosphorylation of STAT proteins.^33, 37-40^ Phosphorylated STAT proteins form homo-or heterodimers and translocate to the nucleus where they bind to consensus sequences in the promoters of target genes^33, 41^. In addition to roles in development and homeostasis, STAT proteins mediate viral defense in NK cells.^42^ Upon IL-2 stimulation, pSTAT5 binds two enhancers located in the 5’ region of *PRF1* promoting its transcription;^43^ upon IL-6 and IL-12 stimulation this enhancer is bound by pSTAT1 and pSTAT4 respectively^44, 45^ STAT5b knockout mice have significantly lower levels of perforin expression at baseline and greatly decreased NK cell cytolytic function.^46^ In humans, STAT5b deficiency is associated with an abnormal NK cell development causing susceptibility to severe viral infections in these patients.^47^ Heterozygous GOF mutations in *STAT1* lead to significantly higher levels of phosphorylated STAT1 (pSTAT1) and increased STAT1 response to type I and II interferons.^48^ These mutations are mostly located in the coiled-coil (CCD) or DNA-binding (DBD) domains and lead to an excess of pSTAT1-driven target gene transcription.^48-50^ Patients with these mutations can develop recurrent or persistent chronic mucocutaneous candidiasis (CMC) or other cutaneous mycosis,^48,49^ staphylococcal infections, disseminated dimorphic fungal infections *(Coccidioides inmitis* and *Histoplasma capsulatum),* viral infections and autoimmune disease.^51-54^

Our investigations of patients with unexplained significant viral susceptibility identified functional NK cell defects in patients with STAT1 GOF mutations, suggesting that STAT1 is important for human NK cell differentiation and function. In this study, we describe an immature, poorly functioning CD56^dim^ NK cell population with low perforin expression and impaired cytotoxic capacity in patients with STAT1 GOF mutations. Administration of the specific JAK1/2 inhibitor ruxolitinib, both *in vitro* and *in vivo,* restored perforin expression in immature CD56^dim^ NK cells and partially restored NK cell cytotoxic function. Together, these data demonstrate effects of ruxolitinib treatment and identify decreased perforin expression and impaired terminal maturation as contributing to functional NK cell defect in STAT1 GOF patients.

## METHODS

### Study approval

Blood samples were collected from patients and healthy donors with consent under approved protocols from Baylor College of Medicine Institutional Review Board, and National Institutes of Health (NIH). All samples were collected with sodium heparin collection tubes.

### Flow cytometry and antibodies

Peripheral blood mononuclear cells (PBMCs) from healthy donors and patients were purified by Ficoll-Paque Plus density gradient centrifugation (GE Healthcare, Pittsburgh, PA). NK cells from all patients with confirmed *STAT1 GOF* mutations were studied phenotypically by FACS and evaluated for NK cell activating, adhesion, inhibitory, and maturation markers as well as intracellular cytokines and lytic granule content. Intracellular cytokines were evaluated in cells stimulated with PMA and Ionomycin (Sigma Aldrich, St. Louis, MO) for 6 hours. Brefeldin A (final concentration 10 ug/mL-Sigma Aldrich, St. Louis, MO) was added 3 hours before antibody staining. The cells were fixed and permeabilized with Cytofix/Cytoperm (BD Biosciences). The antibodies were purchased from BD (CD69, FN50; CD16, B73.1; CD244, 2-69; CD11a, HI111; CD11b, ICRF44; CD18, 6.7; CD94, HP-3D9; Perforin, δG9; CD28, L293), BioLegend (CD56, HCD56; CD3, OKT3; CD16, 3G8; CD8, RPA-T8; NKp46, 9E2; DNAM-1, 11A8; NKG2D, 1D11; CD45, HI30; NKp30, P30-15; CD158b, DX27; CD158d, mAb33; CD62L, DREG-56; CD127, A019D5; CD117, 104D2; CD94, DX22; CD34, 581; GM-CSF, BVD2-1C11; TNF-α, Mab11; IFN-δ, 4S.B3; IL-10, JES3-9D7; IL-13, JES10-5A2), Beckman Coulter (NKp44, Z231; CD25, B1.49.9; CD2, 39C1.5; CD57, NC1; CD122, CF1), eBioscience (CD158a, HP-MA4; CD27, 0323; CD107a, eBioH4A3), R&D Systems (CD159c, 134591; CD159a, 131411; CD215, 151303), and Invitrogen (Granzyme B, GB11). Data was acquired with LSR-Fortessa (BD) cytometer and analyzed using FlowJo (Tree Star, Ashland, OR, USA). NK cell subsets were identified as CD56^bright^CD3^-^ or CD56^dim^CD3^-^. The percentage of NK cells positive for the receptor of interest was defined using a corresponding fluorescent minus one (FMO). For ruxolitinib assays, PBMCs and YTS cell lines were incubated for 48 hours in RPMI supplemented medium with 1000 nM of Ruxolitinib (Selleckchem). After this time the cells were recovered, washed and stained for NK cell receptor expression analysis.

### Cytotoxicity assays

ADCC and NK cell cytotoxicity were measured with Cr^51^ release assay as previously described.^55^ ADCC was evaluated with Raji cell line incubated in the presence or absence of anti-CD20 (Rituximab) (20 μg/mL) and co-cultured with fresh PBMCs for 4 hours at 37°C in 5% CO_2_. For natural cytotoxicity, PBMCs from patients and healthy donors were incubated for 4 hours with IL-2 (1000 U/mL) and the K562 target cell line. YTS and NK92 cell cytotoxicity was evaluated with K562 cell line using a 10:1 effector to target ratio.

### STAT activation assays

STAT1 phosphorylation was measured by flow cytometry after stimulation with IFNα (10 ng/mL-Millipore) for 30, 60, and 120 minutes. STAT5 phosphorylation was measured after stimulation with IL-2 (10 ng/mL, Cell Signaling) for 30. In the final 30 minutes of activation cells were stained with anti-CD3 and anti-CD56 antibodies (Biolegend). After these times the cells were fixed with Fixation Buffer (BD Biosciences) for 10 minutes at 37°, washed with Stain buffer (BD Biosciences) and permeabilized with Perm Buffer III (BD Biosciences) for 30 minutes on ice. Cells were stained with pY701-STAT1 AlexaFluor 488, and pY694-STAT5 Alexa Fluor 647 (BD Phosflow) antibodies. The samples were collected with LSR Fortessa cytometer (BD) and analyzed with FlowJo software (TreeStar, Ashland, Ore, USA).

### Proliferation assay

Proliferation was evaluated using CellTrace CFSE Cell Proliferation Assay (Life Technologies, Eugene Oregon). PBMCs were stained with 1 μM of CFSE, stimulated with IL-15 (5 ng/mL) and incubated with RPMI supplemented medium at 37°C in 5% CO_2_ for 5 days. The cells were then washed and stained with CD3 and CD56 antibodies (BioLegend). CFSE intensity was measured by flow cytometry and proliferation was evaluated in the CD56^+^CD3^-^ population.

### Cell lines

NK92-dependent interleukin-2 (IL-2) Natural killer cell line is derived from rapidly progressive non-Hodgkin’s lymphoma cells, these cells expresses the following surface marker CD2, CD7, CD11a, CD28, CD45, CD54, this cell line not express CD1, CD3, CD4, CD5, CD8, CD10, CD14, CD16, CD19, CD20, CD23, CD34, and HLA-DR.^56^ NK92 cells were cultured at 37°C in Myelocult H5100 medium (StemCell technologies) supplemented with Penicillin/Streptomycin (Gibco Life technologies) and 100 U/mL IL-2 (Cell Signaling).

The natural killer cell lymphoblastic leukemia (YTS) cell line expressing CD56,^57^ was cultured at 37°C in RPMI 1640 (Gibco Life Technologies) supplemented with Penicillin/Streptomycin (Gibco Life Technologies), non-essential amino acids (Gibco Life Technologies), 1mM Sodium Pyruvate (Corning Cellgro), 1M HEPES solution (Gibco Life Technologies), and L-Glutamine (Gibco Life Technologies). STAT1 mutant NK cell lines were generated by CRISPR-Cas9 gene editing and performed by nucleofection of 5 ug of GeneArt CRISPR Nuclease Vector plasmid containing the guide sequences ATCACTCTTTGCCACACCATGTTT, ACTGTTGGTGAAATTGCAAGTTTT, and TTCAGCCGCCAGACTGCCATGTTTT (Life Technologies) using the Amaxa nucleofector and Lonza-Kit R. 24 hours post-nucleofection GFP positive cells were sorted. Single clones were expanded and STAT1GOF positive cells were confirmed by flow cytometry.

### Statistics

The statistical significance of differences between healthy donors and patients was determined by unpaired two-tailed Student’s t-test. Differences between multiple groups were determined by Ordinary one-way ANOVA test. Statistical analyses were conducted using Prism 7.0 (GraphPad Software).

## RESULTS

### Patients and clinical manifestations

We evaluated 16 patients with STAT1 GOF disease. The male to female ratio was 1:1, and median age was 18.18 years (range: 3-31 years). Two of these patients died from progressive fungal infections. The clinical features are listed in Table 1. *Immune manifestations*. Two patients had IgA deficiency (<7mg/dL). IgG deficiency varied; two patients received IgG replacement. CD4 lymphopenia was present in 3 patients. Eleven patients had <0.5% CD4+IL-17+ cells (normal range 0.5-2.5%).

### STAT1 mutations

Sequencing of *STAT1* identified heterozygous mutations in each patient. In 8 patients, the mutations were in the coiled-coil domain (CCD) and the remainders were in the DNA-binding domain (DBD) (Table I). Out of eight discrete mutations, seven have been previously reported;^51, 53, 54^ c.983A>G p.H328R is novel.

**Table I.** Clinical manifestations and STAT1 mutations. CMC, Chronic Mucocutaneous Candidiasis; CMV, Cytomegalovirus; HSV, Herpes simplex virus; IPEX, <u>I</u>mmune dysregulation, <u>P</u>olyendocrinopathy, <u>E</u>nteropathy, <u>X</u>-linked; VZV, Varicella zoster virus; TB, *Tubercle bacillus:* M, male; F, Female; R, patient with ruxolitinib treatment; +, died.

| Patient<br>Gender/Age (y) | Mutation | CMC | Other clinical manifestations |
| --- | --- | --- | --- |
| **Coiled-coil domain mutations** |  |  |  |
| Patient 1<br>F/15 | c.606G>A<br>p.M202I | CMC | TB pneumonia<br>Cryptococcal pneumonia |
| Patient 2<br>M/3 | c.820C>T<br>p.R274W | CMC | Influenza infection<br>Adenopathy |
| Patient 3<sup>R</sup><br>F/31 | c.820C>T<br>p.R274W | CMC | Papillary thyroid CA<br>Skin squamous cell CAs<br>Diabetes Mellitus type I<br>Recurrent oral/vaginal aphthous ulcers<br>Sinopulmonary bacterial infections<br>Skin and soft tissue infections<br>Hypertension |
| Patient 4<br>F/10 | c.820C>T<br>p.R274W | CMC | Thrush<br>Growth failure<br>Vulvar erythema<br>Mild erythematous facial rash<br>Persistent vaginal discharge |
| Patient 5<br>M/5 | c.853C>A<br>p.Q285K | CMC | Growth failure<br>*M. fortuitum* lymphadenitis<br>Sinopulmonary bacterial infections<br>Skin and soft tissue bacterial infections<br>Rectal lesions |
| Patient 6<br>M/5 | c.866A>G<br>p.Y289C | CMC | CMV viremia<br>HSV gingivostomatitis<br>Recurrent thrush<br>*P. jirovecii* pneumonia |
| Patient 7<br>M/8 | c.866A>G<br>p.Y289C | None | None |
| Patient 8<br>F/30 | c.866A>G<br>p.Y289C | CMC | None |
| **DNA-binding domain mutations** |  |  |  |
| Patient 9<sup>R</sup><br>M/7 | c.983A>G<br>p.H328R | None | IPEX-like enteropathy<br>Diabetes mellitus type I<br>Hypothyroidism<br>Growth failure<br>*M. fortuitum* lymphadenitis<br>Chronic sinopulmonary infections<br>Chronic norovirus enteritis |
| Patient 10<br>F/21 | c.986C>T<br>p.P329L | CMC | HSV stomatitis |
| Patient 11<sup>+,R</sup><br>F/22 | c.1057G>A<br>p.E353K | None | Oral herpes<br>Osteomyelitis with spinal cord lesion by coccidioidomycosis disseminated<br>Persistent tinea capitis and kerion by *T. tonsurans* |
| Patient 12<br>F/27 | c.1057G>A<br>p.E353K | CMC | *T. tonsurans* hand granuloma |
| Patient 13<br>M/15 | c.1154C>T<br>p.T385M | CMC | Selective IgA deficiency<br>Recurrent sinopulmonary disease<br>Bronchiectasis<br>*S. aureus* and *S. pneumoniae* infections |
| Patient 14<sup>+</sup><br>F/16 | c.1154C>T<br>p.T385M | CMC | IPEX-like Enteropathy<br>Herpes zoster infection<br>VZV infection<br>CMV infections<br>Mild eczema<br>Hepatosplenomegaly<br>Bronchiectasis<br>Generalized osteopenia<br>Aneurysms |
| Patient 15<br>M/28 | c.1154C>T<br>p.T385M | CMC | VZV infections<br>Histoplasmosis<br>Bronchiectasis<br>Single cervical node with *M. fortuitum* |
| Patient 16<br>M/30 | c.1154C>T<br>p.T385M | CMC | Recurrent HSV infections<br>Molluscum contagiosum<br>Enteropathy<br>Sinopulmonary bacterial infections<br>Bronchiectasis<br>Diabetes mellitus type I |

*In vitro* STAT1 activation was tested in primary patient NK cells representing 5 mutations (3 CCD and 2 DBD) in response to IFN-α. We observed persistent phosphorylation without appropriate dephosphorylation in all patients NK cells when compared to healthy donors (Fig 1, A). Furthermore, mutations in the DBD had significantly higher levels of activated STAT1 after 120 minutes of stimulation than mutations located in the CCD (Fig 1, B). Interestingly, we observed that patients with mutations in the DBD tend to have a more severe clinical phenotype than patients with CCD mutations (Table I).

**FIG 1.**
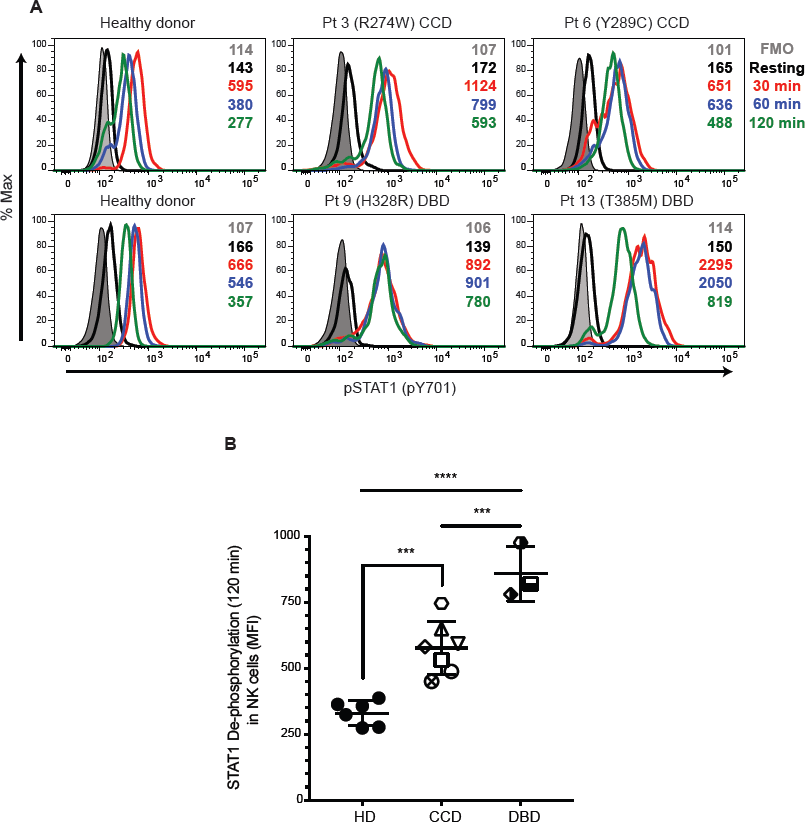
STAT1 GOF mutations lead to delayed dephosphorylation in NK cells. Dephosphorylation in NK cells stimulated with IFN-α for the indicated periods in healthy donors, **A**, patients with mutations into the Coiled-coil domain (CCD; patients 2, 3, 4, 5, 6, 7, and 8) or DNA-binding domain (DBD; patients 9, 13, and 15). **B**, levels of STAT1-activated after 120 minutes of stimulation. HD, healthy donor; Pt; patient. The statistical significance was analyzed using the Ordinary one-way ANOVA test \*\*\**p*<0.001, \*\*\*\**p*<0.0001.

### Patients with STAT1 GOF mutations have decreased NK cell cytotoxic function

We performed standard Cr^51^ release assays using PBMCs from 14 patients and healthy donor controls using K562 target cells in the presence and absence of exogenous IL-2. We compared the NK cell cytotoxicity activity from each patient with a representative NK cell cytotoxicity from a pool of 10 healthy donors. NK cells from most patients with CCD mutations had residual cytotoxic function that was partially responsive to IL-2 (Fig 2, A; and see Fig E1, A,). In contrast, cells carrying mutations in the DBD had more severe defects in their NK cell cytotoxicity that was not improved by IL-2 (Fig 2, B; and see Fig E1, B). NK cells from 10 patients were incubated with Rituximab-opsonized Raji target cells to assay antibody-dependent cellular cytotoxicity (ADCC). All patients with CCD mutations had residual ADCC (see Fig E2, A,), whereas, ADCC in cells from patients with DBD mutations was more impaired (see Fig E2, B). These data suggest that impairment of NK cell function is affected by the location of the *STAT1* mutation.

**FIG 2.**
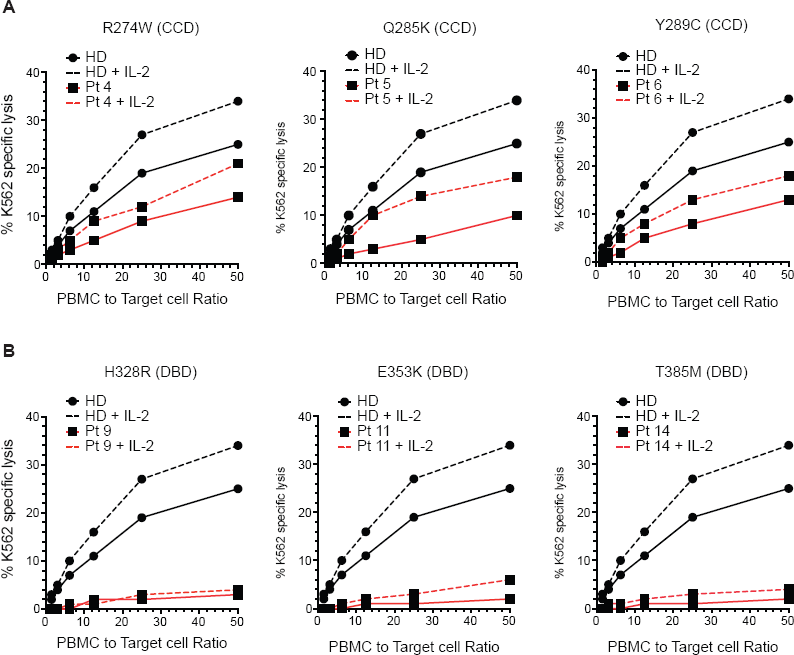
Impaired NK cell cytotoxic capacity in patients with *STAT1 GOF* mutations. NK cell cytotoxicity was measured by Cr^51^ release in the presence or absence of 1000 U/mL of IL-2. **A**, patients with Coiled-coil domain (CCD) mutations; **B**, patients with DNA-binding domain (DBD) mutations. HD, healthy donor; Pt; patient.

### Impaired terminal maturation of the CD56^dim^ NK cell subset in patients with *STAT1 GOF* mutations

NK cell targeted lysis is mediated and enabled by activating and inhibitory receptors on mature NK cells. To determine if the impaired NK cell cytotoxic function found in our patients was associated with a demonstrable NK cell phenotype, we performed flow cytometry for NK cell developmental markers, functional NK cell receptors (activating and inhibitory), intracellular cytokines, and lytic granule contents^7, 10, 25, 58^. NK cell subsets were identified as CD56^bright^CD3^-^ or CD56^dim^CD3^-^ (Fig 3, A). We observed lower NK cell frequencies in STAT1 GOF patients (4.0±2.5%) (mean±SD) compared to healthy donors (9.7 ± 2.1%). The frequency of CD56^bright^ NK cell subset in peripheral blood from patient and control groups was similar. However, patients with *STAT1 GOF* mutations had a decreased frequency of cells within the CD56^dim^ NK cell subset (2.631±1.774%) in peripheral blood than healthy donors (9.894%, range 7.25-13.7%) (Fig 3, B), suggesting impaired terminal maturation or survival of the CD56^dim^ NK cell subset.

In healthy donors, the CD56^dim^ NK cell subset is characterized by high levels of expression of CD16 and perforin and acquisition of CD57 and CD8.^7, 15^ Patients with *STAT1 GOF* mutations have abnormal expression with low mean fluorescence intensity (MFI) of CD16, perforin, CD57, CD8 (Fig 3, C; and see Fig E3, A,). STAT1 GOF patients had decreased_percentage of CD56^dim^perforin^+^ NK cells (70.94 ± 23.4%) and CD56^dim^CD16^+^ NK cells (61.03± 13.62%) (see Fig E3, B) compared to healthy donors ((91.44 ± 3.602%), (95.55 ± 3.588%)) respectively. Patient NK cells also had higher expression of receptors associated with the CD56^bright^ subset, including CD94, CD117, CD94, and NKG2A (Fig 3, D and E3, B). While the CD56^bright^ subset in NK cells from patients with *STAT1 GOF* mutations had higher levels of granzyme B and lower levels of TNF-α (see Fig E3, C), the CD56^dim^ subset appears to have impaired terminal maturation. These findings suggest that the CD56^dim^ subset in STAT1 GOF patients is immature and potentially non-functional.

**FIG 3.**
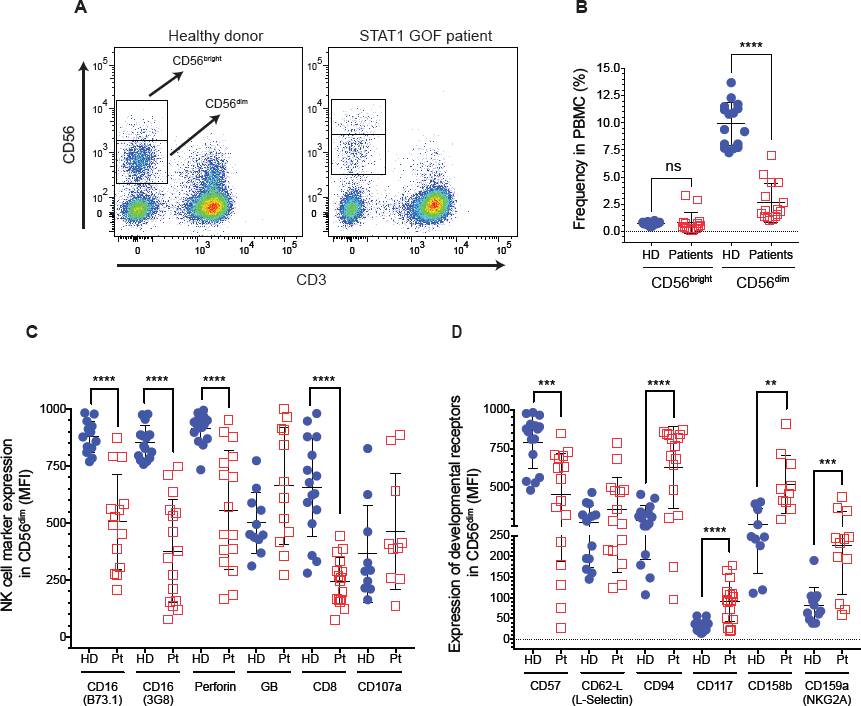
CD56^dim^ NK cells have an immature phenotype in patients with *STAT1 GOF* mutations. NK cell phenotype from healthy donors and STAT1 GOF patients were analyzed by flow cytometry. **A**, NK cell subsets were defined as CD56^bright^CD3^-^ cells or CD56^dim^CD3^-^. **B**, representative graph shows the frequencies of CD56^bright^CD3^-^ and CD56^dim^CD3^-^ NK cells subsets. **C**, representative graphs shows mean fluorescence intensity (MFI) for each NK cell receptor expressed by CD56^dim^ NK cells. Each blue circle represents a healthy donor and a red box represents a STAT1 GOF patient. Horizontal bars represent means and the vertical bars indicate the standard deviation. The statistical significance was analyzed using the unpaired Student’s t-test. Results are representative of 3 independent experiments. \*\**p*<0.01, \*\*\**p*<0.001, \*\*\*\**p*<0.0001.

### STAT1 GOF mutations lead to decreased perforin expression and cytotoxic capacity in NK cell lines

To further delineate the effect of STAT1 dysregulation on NK cell phenotype and function, we modeled two of the DBD mutations (E353K and T385M) in YTS NK cell lines using CRISPR-Cas9 mediated gene editing. While expression of total STAT1 was not affected by either mutation in either cell line (data not shown), both mutant cell lines showed enhanced STAT1 phosphorylation in response to IFN-α stimulation compared with WT cells (Fig 4, A). To further elucidate the abnormality seen in the primary cells, we looked at select developmental markers in this cell line. The expression of CD94 and NKG2A was increased in the mutant cell lines compared to WT (Fig 4, B). Furthermore, we detected decreased perforin expression of YTS STAT1 mutant cell lines compared to WT (Fig 4, B). To determine whether these phenotypic differences corresponded with an effect on function, we performed Cr^51^ release cytotoxicity assays and observed that the cytotoxic capacity of YTS STAT1 mutants was decreased more than 4-fold compared with the WT (Fig 4, C). Similarly, expression of the CCD M202I mutation in a different NK cell line, NK92, led to an analogous functional phenotype (Fig 4, D and see Fig E4) with decreased perforin expression and reduced NK cell cytotoxic function (Fig 4, D). This phenotype was consistent with our results from primary patient cells, as the effect of CCD mutations was milder than that of the DBD mutations and define the impact of patient-derived mutant STAT1 GOF on NK cell perforin expression and maturation signatures.

**FIG 4.**
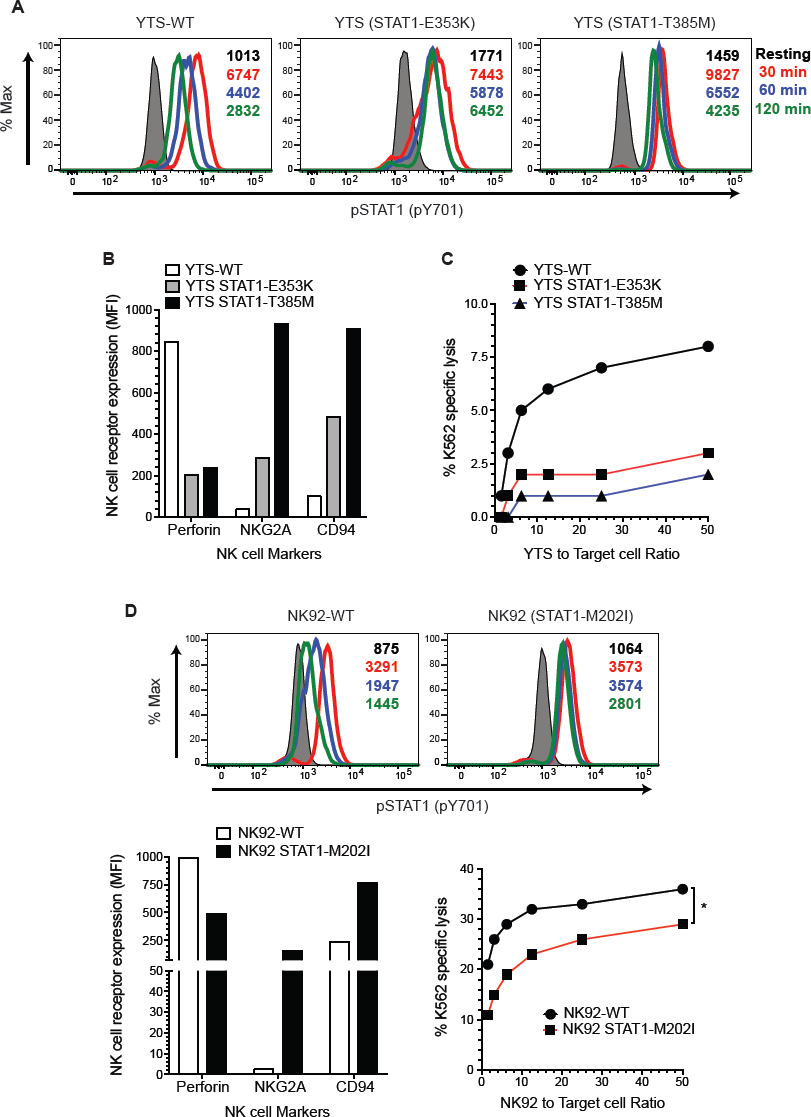
*STAT1 GOF* mutations affect the cytotoxic capacity and perforin expression in NK cell lines. DNA-binding domain mutations E353K and T385M were expressed in the YTS cell line using CRISPR-Cas9 gene editing. **A**, STAT1 hyper-phosphorylation in response to IFN-α was measured by intracellular flow cytometry using pY701-STAT1 antibody. **B**, the phenotype of YTS STAT1 GOF cell lines was measured by flow cytometry. **C**, NK cell cytotoxicity was analyzed against K562 target cells using standard Cr^51^ release protocol. **D**, STAT1 hyper¬phosphorylation in response to IFN-α; perforin, NKG2A, and CD94 expression, and cytotoxic capacity measured in NK92 STAT1-M202I mutant cell line. MFI, Mean fluorescence intensity; WT, wild type. Results are representative of 3 independent experiments. * *p*<0.05.

### STAT1 GOF mutations impair STAT5 phosphorylation in NK cells

The functional assays were performed in patients with mutations in the CCD (patients 2, 4, 5, 6, 7, and 8) and DBD (patients 11, 15, and 16). Considering the vital STAT-related role of IL-15 in NK cell development and homeostasis, we evaluated the proliferative capacity of NK cells from STAT1 GOF patients in response to IL-15 stimulation. All patients tested had significantly decreased proliferation compared to healthy donors, independent of the affected domain (Fig 5, A).

**FIG 5.**
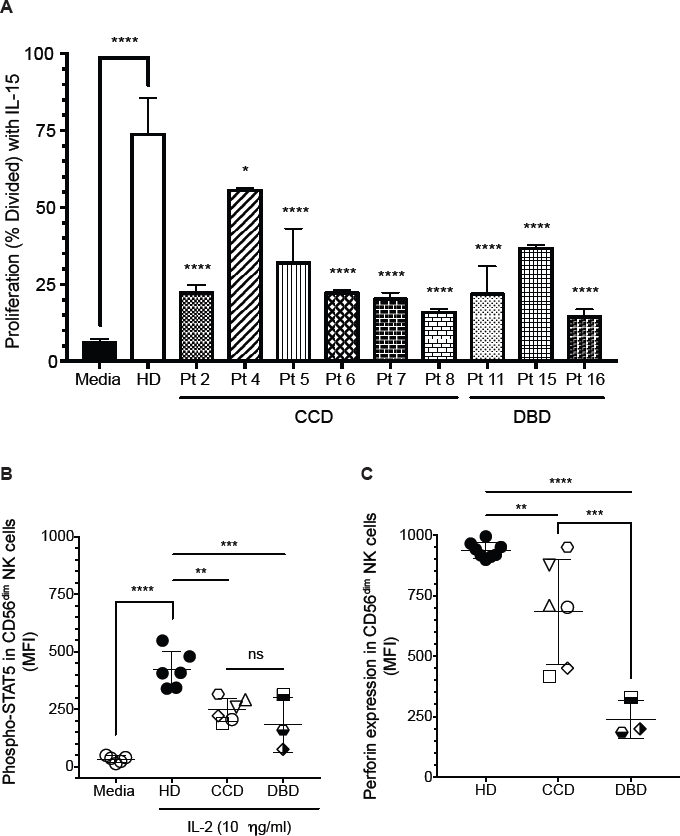
Decreased STAT5 activation in CD56^dim^ NK cells from STAT1 GOF patients. **A**, NK cell proliferation from healthy donors and STAT1 GOF patients in presence of IL-15 (5 ng/mL). **B**, phospho-STAT5 levels in CD56^dim^ NK cells stimulated with IL-2 (10 ng/ml) for 30 minutes. Each point represents the mean fluorescence intensity for each individual healthy donor or patients. **C**, perforin expression levels in CD56^dim^ NK cells from patients with CCD or DBD mutations. Statistical significance was analyzed using the Ordinary one-way ANOVA *test.* * *p*<0.05, **p<0.01, ***p<0.001, ****p<0.0001. HD, healthy donors; MFI, Mean fluorescence intensity; CCD, Coiled-coil domain (patients 2, 4, 5, 6, 7, and 8); DBD,DNA-binding domain (patients 11, 15, and 16). Results are representative of 2 independent experiments for each patient.

Significant crosstalk between STAT signaling molecules exists, including antagonistic binding to common enhancer elements. In particular, the competitive binding of STAT3 and STAT5 to a common 5’ perforin enhancer has been demonstrated in T lymphocytes, dendritic cells, and cancer cell lines.^18,43,59^ On other hand, STAT5 induces perforin expression in response to IL-2.^43^ To determine whether the delayed dephosphorylation of STAT1 affects the activation of STAT5 and leads to the significantly decreased perforin expression in NK cells from patients with *STAT1 GOF* mutations, we evaluated STAT5 activation after IL-2 stimulation. We observed significantly decreased STAT5 activation in response to IL-2 in CD56^dim^ NK cells from the CCD and DBD mutations (Fig 5, B). Interestingly, patients with DBD mutations had lower levels of perforin compared with CCD patients (Fig 5, C). The lower levels of phosphorylated STAT5 in DBD patients after 30 minutes of stimulation corresponds with the degree of impairment of perforin expression in STAT1 GOF NK cells (Fig 5, B and C).

To determine whether low NK cell proliferation with IL-15 stimulation and low STAT5 activation in response to IL-2 was due to abnormal IL-2 and IL-15 receptor expression, we measured their expression by flow cytometry. We found normal receptor expression for IL-2 and IL-15 in healthy donors and affected subjects (data not shown). Therefore, the degree of impairment of NK cell proliferation and perforin expression in STAT1 GOF NK cells tracks with the degree of impairment in STAT5 activation.

### Ruxolitinib treatment restores perforin expression and cytotoxicity in CD56^dim^ NK cells from STAT1 GOF patients

Recently, three studies reported the efficacy of JAK1/2 pathway inhibition by ruxolitinib in patients with STAT1 GOF mutations. Constitutive hyper-phosphorylation of STAT1 in these patients was inhibited while STAT3 phosphorylation was not affected. Importantly, ruxolitinib treatment led to improvement of CMC and autoimmunity.^60-62^ We sought to determine whether ruxolitinib affected the phenotype of NK cells from patients with STAT1 GOF mutations. We evaluated the NK cell phenotype and function from three patients (3, 9, and 11) before and during oral ruxolitinb treatment. We observed higher levels of CD56^dim^Perforin^+^ NK cells after treatment with restoration of perforin expression on CD56^dim^ NK cells during ruxolitinib treatment (Fig 6, A and see Fig E5, A), accompanied by a 5-fold increase in NK cytotoxicity in response to IL-2 in patient 9 and a 2-fold increase in patient 11 (Fig 6, B). In addition, the ADCC from patient 11 showed a 2-fold increase (see Fig E5, C). Increased perforin expression accompanied improved NK cytotoxic function. We did not observe an increase in NK cytotoxicity in patient 3 likely do to the short duration of treatment when the assay was performed (Fig E5, B). Patient 9 and 11 had been on ruxolitinib for at least 3 months when the post-ruxolitinib assays were performed, whereas, Patient 3 was treated for only 2 weeks. Perforin restoration is immediate while improvement in cytotoxicity comes with a longer duration of therapy. Weinacht et al^62^ reported a patient with clinical and laboratory improvement after a year on ruxolitinib treatment. Taking our experiences together, it appears at least 3 months of therapy with Ruxolitinib is needed to demonstrate improvement in function.

**FIG 6.**
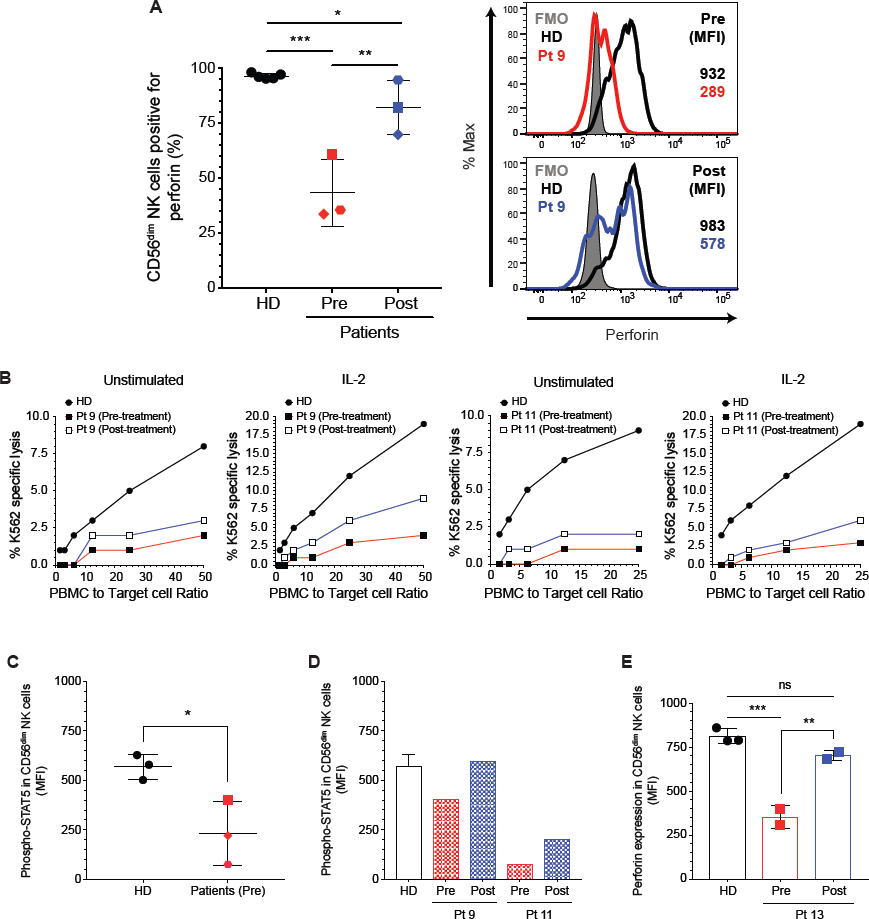
Ruxolitinib restores perforin expression in primary NK cells. **A**, percentages of CD56^dim^Perforin^+^ NK cells and perforin expression on CD56^dim^ NK cell subset pre-and post-treatment. **B**, natural cytotoxicity was performed with Cr^51^ release protocol with freshly isolated PBMC from patient 9, and 11 and a respective healthy donor and was tested against K562 target cells in the presence or absence of 1000 U/mL of IL-2. STAT5 activation in CD56^dim^ NK cells after stimulation with IL-2, **D**, patients 3, 9, and 11 (pre-treatment) and **D**, in CD56^dim^ NK cells from patient 9 and 11post-treatment. **E**, PBMCs from Pt 13 were incubated *in vitro* for 48 hours with or without 1 M of ruxolitinib and the perforin expression was measured in CD56^dim^ NK cells. The statistical significance was analyzed using the Ordinary one-way ANOVA test * *p*<0.05, \*\**p*<0.01, \*\*\**p*<0.001.

Interestingly, while the expression of perforin increased, CD16, CD8, and CD57 expression was not affected (see Fig E5, D). Despite the dramatic increase in perforin and improved NK cell cytotoxicity, the course of the fungal infection was so advanced in patient 11 when treatment was started that her disease was not ameliorated and she eventually died. However, on acyclovir prophylaxis, she did not have any breakthrough herpes infections. Patient 9 experienced significant improvement in his enteropathy and chronic norovirus. His stool output decreased from 3 liters of stool per day to 2-3 loose stools 4 weeks after treatment began, and then 1-2 formed stools per day by 3 months post treatment.

To determine whether ruxolitinib had a direct effect on perforin expression, we evaluated the STAT5 phosphorylation after stimulation with IL-2 in CD56^dim^ NK cells. Our findings showed a lower level of STAT5 phosphorylation in CD56^dim^ NK cells with IL-2 stimulation (Fig 6, C), which also corresponds with low perforin expression in pre-treatment samples. After ruxolitinib treatment, we observed higher levels of STAT5 activation in post-treatment sample from patient 9 and 11 (Fig 6, D). These results demonstrate that decreased activation of STAT1 through JAK inhibition affects the activation of STAT5 and, in turn, perforin expression. To compliment the studies on the *in vivo* effects of ruxolitinib treatment over perforin expression, we incubated PBMCs isolated from another patient (Pt 13) for 48 hours with 1 μM of ruxolitinib. Perforin expression was partially restored in CD56^dim^ NK cells (Fig 6, E); whereas, the levels of perforin observed in CD56^dim^ NK cells from healthy donors were not affected. Similar results were observed using the STAT1 GOF NK cell line (see Fig E5, E). Thus, inhibition of JAK1/2 signaling *in vivo* and *in vitro* leads toward normalized STAT1 phosphorylation and restoration of NK cell function in STAT1 GOF NK cells.

While the maturation defect in STAT1GOF NK cells is not rescued, direct modulation of STAT1 phosphorylation with ruxolitinib can partially remediate poor NK cell function in these patients.

## DISCUSSION

Patients with *STAT1 GOF* mutations have impaired IL-17A and IL-17F immunity, associated with CMC.^48, 49, 53^ However, impaired IL-17 immunity does not fully explain the susceptibility to viral infections.^52, 54^ *STAT1* loss-of-function mutations have been reported with impaired NK cell activity and CMV infection.^31^ The association of herpesvirus infection with NK cell deficiency led us to investigate the phenotype and function of NK cells in patients with *STAT1 GOF* mutations. We have identified that excess STAT1 phosphorylation owing to these mutations is associated with impaired NK cell terminal maturation, decreased expression of the critical cytolytic effector molecule perforin, and impaired cytolytic function. Importantly, these effects can be partially ameliorated *in vivo* and *in vitro* by treatment with the JAK1/2 inhibitor ruxolitinib.

We found that the STAT1 GOF-associated CD56^dim^ NK cell subset have an immature phenotype, characterized by aberrant expression of CD94, CD117, NKG2A, molecules associated with the CD56^bright^ subset, as well as low levels of CD16, perforin, CD57 and CD8. While a similar developmentally intermediate population typically makes up between 20% and 40% of CD56^dim^ NK cells in healthy donors,^11, 19, 63^ it was the majority (80%) of CD56^dim^ NK cells found within the peripheral blood of STAT1 GOF patients. Notably, The STAT1 GOF CD56^dim^ population is characterized by low levels of CD16 and perforin; two important components of the cytotoxic capacity to eliminate infected cells.

Interestingly, DNA-binding domain mutations generally conferred a more profound NK cell defect than those in the coiled-coil domain. The *STAT1* domain mutated is important in the severity of the NK cell dysfunction in STAT1 GOF patients. However, expression of both the coiled-coil and DNA-binding domain mutations in NK cell lines equally affected perforin expression and cytotoxic capacity, suggesting that additional factors may dictate the effect of *STAT1* mutation on NK cell function. This variability in phenotype has also recently been reported by Tabellini et al., who describe a cohort of STAT1 GOF patients with normal perforin and CD16 expression.^32^

STAT5 phosphorylation and signaling, including IL-15 mediated proliferation, are impaired in STAT1 GOF. STAT5’s unique role in NK cell proliferation and cytotoxic activity suggests that its impairment may cause at least some of the NK cell maturation defect seen in STAT1 GOF patients.^18, 46^ In addition, the binding of *STAT5* to the 5’ enhancer in the *PRF* gene likely contributes specifically to the down-regulation of perforin in patient NK cells. The re-expression of perforin upon reduction of STAT1 phosphorylation suggests that this is not a hardwired developmental defect. The mechanism of suppression of STAT5 by excess STAT1 phosphorylation is unclear. STAT1-induced SOCS-mediated negative regulation of STAT5 is one possibility, and the recent description of increased SOCS1 and SOCS3 expression in STAT1 GOF patient NK cells supports this hypothesis.^32^ The lack of rescue of overexpression of CD94/NKG2A and CD117 on CD56^dim^ NK cells following ruxolitinib treatment suggests that these markers are not directly modulated by excess STAT1 signaling.

With the onset of genetic testing in primary immunodeficiency disease(PID), we can now choose therapies that modulate a specific target.^64^ For example, abatacept is now successfully used to treat CTLA-4 haplo-insufficiency as well as other directed cytokine and biologic therapies for a wide range of PIDs. ^65^ Autoimmune disease in STAT3 GOF disease was successfully treated with IL-6R blockade.^66^ Treatment with the JAK 1/2 inhibitor ruxolitinib in a patient with refractory candida esophagitis led to resolution of many symptoms within a 5-month period.^60^ Increases in IL-17 production in STAT1 GOF patients following treatment with ruxolitinib was also associated with reduced burden of CMC, reduced hyperresponsiveness to type I and II interferons, normalized T_H_1 and follicular T helper responses, and had remission of the autoimmune mediated cytopenias.^61, 62^ Our results demonstrate that perforin expression and NK cell cytotoxic capacity are partially restored following *in vivo* treatment, suggesting that ruxolitinib may improve NK cell function, but its effect on other lymphoid and myeloid elements must be assessed to determine the therapeutic utility of ruxolitinib in STAT1 GOF disease.

In conclusion, STAT1 GOF mutations cause a distinct NK cell phenotype likely owing to an impairment of NK cell terminal maturation. These mutations cause poor perforin expression and this NK cell population has decreased or absent NK cell cytotoxic activity. Importantly, perforin expression and NK cell function are restored by JAK inhibition with ruxolitinib. These data indicate that STAT1 has critical roles in NK cell development and maturation, through direct mediation of cytotoxic mediators and through modulation of other mediators, including STAT5.

## Online Repository figures

**FIG E1.** Natural cytotoxicity was measured in the presence or absence of 1000 U/mL of IL-2. **A**, patients with CCD mutations; **B**, patients with DBD mutations. HD, healthy donor; Pt; patient.

**FIG E2.** Antibody-dependent cellular cytotoxicity (ADCC) was performed using Raji target cells in the presence or absence of Rituximab (20 ug/mL) in patients with **A**, CCD and **B**, DBD mutations. HD, healthy donor; Pt; patient.

**FIG E3.** NK cell subsets phenotype in patients with *STAT1 GOF* mutations. **A**, representative perforin expression plots in total NK cells. Positive **B**, CD56^dim^ or **C**, CD56^bright^ NK cells for each receptor. Each blue circle represents a healthy donor and a red box represents a STAT1 GOF patient. Horizontal bars represent means and the vertical bars indicate the standard deviation. The statistical significance was analyzed using the unpaired Student’s t-test. Results are representative of 3 independent experiments. * *p*<0.05, ** *p*<0.01, *** *p*<0.001, **** *p*<0.0001.

**FIG E4.** NK cell receptors expression in NK92 STAT1-M202I cell line. The expression was measured by flow cytometry, the expression was compared with WT cell line and respective fluorescence minus one (FMO). Each histogram showed the FMI for each cell line.

**FIG E5.** Ruxolitinib restore the perforin expression in STAT1 GOF NK cell lines. **A**, perforin expression in CD56^dim^ NK cell from patient 3 pre-and post-treatment. **B**, natural cytotoxicity from patient 3 and a respective healthy donor in the presence or absence of 1000 U/mL of IL-2. **C**, ADCC from patient 11 performed using Raji target cells in the presence or absence of Rituximab (20 ug/mL) before and during her treatment. **D**, CD56^dim^ phenotype in patients with oral ruxolitinib treatment, **E**, perforin expression in YTS STAT1 GOF NK cell lines after 48 hrs of stimulation with 1 μM of ruxolitinib. Statistical significance was analyzed using the Ordinary one-way ANOVA *test*. \**p*<0.05, \*\**p*<0.01, \*\*\*\**p*<0.0001.

